# Twelve Elements of Visualization and Analysis for Tertiary and Quaternary Structure of Biological Molecules

**DOI:** 10.1101/153528

**Authors:** Philippe Youkharibache

## Abstract

During the last decades, 3D Molecular Graphics in Life Sciences has been used almost exclusively by experts through complex software and applications ranging from Structural Biology to Computer Aided Drug Design. The emergence of JavaScript and WebGL as a viable platform has enabled 3D visualization of biomolecular structures through Web browsers, without any need for specialized software. Although still in its infancy, Web Molecular Graphics opens new perspectives. This white paper, proposes a set of Twelve Elements to consider to enable 3D visualization and structural analyses of biological systems in Web molecular viewers. The Elements go beyond 3D graphics and propose an integrated approach to visualize and analyze molecular entities and their interactions in multiple dimensions, at multiple levels of details, for diverse users. The bridging of 1D sequence browsers and 3D structure viewers, possible under a Web browser, enables information flow where molecular biologists can use structural information directly at the sequence level. Given the tsunami of sequence information linked to diseases from next generation sequencing - in need for interpretation - making structural information readily available to research scientists is a tremendous opportunity for medical discovery. The Twelve Elements are conceptual and are intended to entice developers to architect software components and APIs, and to gather together as a community around common goals and open source software. A few features of emerging viewers, all available as open source, are highlighted. Speed and quality of 3D graphics for large molecular systems, the interoperability of Web components, and the instantaneous sharing of annotated visualizations through the Web, are some of the most amazing and promising capabilities of 3D Web viewing, opening bright perspectives for Life Sciences research.

## Introduction

This white paper is aimed at sharing a few ideas on molecular structure visualization and analysis in the context of biological molecules and their assemblies. It does not intend to be exhaustive. It was originally written early 2015 when I joined NCBI within the NIH while there were discussions about writing a JavaScript 3D viewer to visualize and compare tertiary and quaternary structure of biological molecules and their assemblies, within a Web browser. This was and still is a good move considering progress made in 3D JavaScript and mobile technology.

Rewriting a 3D molecular viewer, inevitably, leads to discussions about **what** should be visualized, **how**, and **why**, but also **who** is the user, or rather who are the various uses and users, from casual to so-called power-users. The irony in the discussions on usage and users is that one can end up losing the perspective on the information **all users need** ultimately and how to convey this information, and make this information not only useful but usable beyond 3D graphics rendering. For some of us, it is déjà vu all over again, and in this case, I hope past experiences can help enrich new developments. This was my sentiment and I thought any developer could benefit from past experiences and could try to get a vision on how far visualization can go. Not visualization for the sake of it but for a purpose: understanding biological macromolecular structure, identifying key molecular interactions, extracting that information so that it can be used for downstream analyses.

Rather than describing past experiences I thought I could condense some of the lessons learned in a practical set of starting requirements which I believe are of universal need for molecular structure analysis. I ended up structuring this document in “Elements”. I chose that word (plural) by analogy to Euclid’s Elements of Geometry. **“Elements” of a molecular viewer are features that, by themselves, enable many derived features, especially in relation to each other’s**. I placed the emphasis on “systemic accessibility” of molecular information, with integration and interconnection between views and viewers (1D, 2D and 3D). This is also a very important issue: there should not be a divide between Sequence (1D) and Structure (3D) but a bridge. Both are molecular representations. Information belongs to the entities they represent. Information should flow and should be represented in both. This is not only about connecting information and representations, it is also about connecting users, whether a structural biologist, a chemist, or a molecular biologist. Some users live and experiment in a sequence (and experimental) world and some live and think in a 3D structure (and theoretical) world. Aren’t these simply perceptions of the same world, as in *Rumi’s Elephant in the Dark*?

With the explosion of sequence information, the need is even greater today to provide and visualize structural information at the sequence level. This will require a lot of attention in years to come with the need to visualize structural information in sequence browsers/viewers, not just in 3D. Visualization of molecular information is not just 3D. These Elements are conceptual and are intended to entice developers to architect software components and APIs to go from concept to action. It would be about time a community gather together around common goals and open source software.

This white paper was written at the start of NCBI’s efforts towards developing a Web based 3D viewer iCn3D, early 2015. Some descriptions may rely on NCBI’s environment. However, the Elements themselves should be universal, or can be revisited to be universal.

## Recent Developments on 3D molecular viewers

Efforts to develop Web based molecular viewers have started in the last few years in a number of labs. All viewers demonstrate that Javascript with today’s support of component libraries and computer power is definitely a viable option for 3D graphics, most likely the best option available today. The biggest issue for viewers is their extensibility down the road both in terms of architecture and in terms of support. An open source environment seems to be mandatory, and a community should gather together to enable a viable visualization and analysis software environment for the long term to tackle a multitude of projects, themselves evolving.

Recently a few developers gathered in Shonan, Japan to discuss common needs on Web molecular graphics. This represents a step towards initiating a community: http://shonan.nii.ac.jp/shonan/blog/2015/10/30/web-%E2%80%90based-molecular-graphics/. Most members who gathered participate to new Web viewers developments. A few examples include NCBI’s **iCn3D** viewer developed by Jiyao Wang (Wang 2017), RCSB’s **NGL** viewer by Alexander Rose(Rose 2015, 2016) or **Jolecule** by Bosco Ho, to name a few. All viewers are available in GitHub as open source (https://github.com/ncbi/icn3d; https://github.com/arose/ngl; https://github.com/boscoh/jolecule), all have specific strengths, due to the emphasis and focus of their developers. What seems important for all these viewers going forward is *sustainability* and *extensibility*.

Looking ahead, we need a flexible and modular architecture, moving towards componentization, developing libraries and APIs so that new best of breed applications can be built quickly. Currently we can already experience a spectrum of capabilities rather unique to each viewer. One has an emphasis on analytical molecular representations, one on rendering speed for viewing and managing very large molecular systems, and one in assisted/guided visualization of molecular features. Componentization has started in NGL and also a viewer supported by PDBe in Europe: **Litemol** by David Sehnal (https://github.com/dsehnal/LiteMol). It is interesting to note that PDBe and the EBI are moving towards developing component libraries *systematically*, not just for 3D graphics (http://www.ebi.ac.uk/pdbe/pdb-component-library/) and this will turn out to be very important to “mashup” data from various sources. It will take an entire community however to develop all the needed sharable component libraries.

Since the core of this document was written in 2015 it makes reference to Cn3D and its applications. This was and still is, NCBI’s original sequence/structure aligner and viewer (Wang 2000 - https://www.ncbi.nlm.nih.gov/Structure/CN3D/cn3d.shtml). I added at the end of the document some links to current web viewers demonstrating today some of the twelve Elements, a work in progress.

## Biological molecular Structure and structural description

The title of this paragraph says it all. It is about Biology. Molecular representations should be able to describe interacting biological molecules. Viewing biological molecules - *and their properties* - should teach us the structural basis of biological function, as well as structure/sequence/function conservation in evolution, since we can also compare structurally molecular assemblies from diverse species.

**Element #1: Allow naming of molecular entities and description of all hierarchical levels from supra-molecular assemblies down to atoms.**

*The system should be able to parse an “atom/molecular spec”, with commonly used nomenclatures for molecular assemblies, molecules, chains, domains, residues, atoms.*

**Element #2: Allow the import of molecular structure coordinates and topologies in widely used formats, generate covalent bonds for organic and biological molecules and visualize molecules in 3D.**

*Direct Interfaces to databases with quality control is the ultimate. In that case, software should be extensible/scalable with database growth.*

One can get at the description of residues/atoms today when picking residues in Cn3D (and also in the new JavaScript viewer), but one cannot select an atom, a residue a list/series of residues by typing its definition, and eventually find it and zoom in. One can get at a Prosite “pattern” in terms of residues, a very useful feature that could be generalized for selection. Picking structures (domains) is also available, but not subsets as they cannot be specified at the user interface level. Some generalization is needed. It is important to be able to specify any part of any assembly, molecule, domain, etc. for display and/or to define a set (see later) or sets of atoms, residues, domains, molecules for calculating properties. For example A:1-5;15-20:CA, or using names *:PRO,LYS: with 3 or 1 letter codes, etc.

## Molecular sets and subsets

The ability to name/label individual or sets of molecules, residues or groups/atoms is, after the introduction of a flexible description language for atoms, molecules and assemblies (atom/molecule spec), one of the most important and fruitful element in opening up analyses. Sets are already available, as defined by computations such as VAST. In Cn3D sets are defined by alignments and can be named and used mostly for coloring. The concept simply needs to be extended to different methods and should be used across the system, and in many cases, sets can be defined at the system level.

**Element #3: Allow flexible selection of any part of molecular entities through subsetting of such entities at any hierarchical level as well as across molecular boundaries, based on topological, geometrical or relational properties.**

*Provide widely used default sets such as secondary structure elements, domains, known biological elements (e.g. PIP box) and active sites.*

*Named Subsets can be seen as annotations as much as structural elements for analysis.*

A number of native sets can and should be available, such as strands and helices (A, B,C … for each SSE) and turns/loops/linkers scan inherit names such as AB, BC, CD etc …. However, it would also be useful for users to introduce their personal sets and nomenclature, should they wish to do so, and override the system’s definition for their particular use, if needed. If this is allowed then they should be able to save this info for future use. This should naturally not affect the NCBI version of information. The simplest way to allow the software to be extensible and customizable is to develop in an open source environment, but ***naming*** should be a system feature.

It should be noted that if a nomenclature is logical and flexible, allowing the representation of any molecular system, the need for a custom nomenclature will be reduced to the ability to define arbitrary sets, name them, and save them. Importantly some sets can be seen as structural elements definitions as well as annotations. A more difficult issue is the introduction system-wide of domain definitions (for discussion). There are also a variety of domain definitions and on my end I use protodomains, i.e. subdomains related through pseudo-symmetry, which are sets and could be handled that way. Introducing sets as dynamic definitions/functions or as specific molecular specs should be discussed.

### Examples of a “Community” or “Sector” Analysis in the literature

We can learn from the field of molecular dynamics, as they monitor multiple molecular variables and properties, many of them are structural, and others are derived/computed from structure. For example, the ability for users to define “communities”, i.e. subsets of structures that crosstalk, reduces complexity in **interaction networks**. In a **community analysis** (McCLendon 2014 and refs cited): one can identify residue-residue correlated motions and partition a protein into structurally contiguous "communities." Most of these communities included 40-60 residues can be associated with a particular protein function or a regulatory mechanism. Well-known motifs based on sequence and secondary structure can often split into different communities, raising the interest for a flexible set selection, as defined earlier. Community maps are sensitive to the presence of different ligands and provide a framework for interpreting long-distance allosteric coupling. See Figure (McCLendon 2014 https://www.ncbi.nlm.nih.gov/pmc/articles/PMC4217441/figure/fig02/

A careful structural analysis with different assemblies/structures under some conditions (e.g.) ligand binding would come to comparable results (Wilson 2015).

The hierarchy of biological structure at the primary, secondary, tertiary, and quaternary levels, is the organization that currently guides our understanding of biological properties and evolutionary origins. Distinct structural organizations can be defined, through a decomposition of proteins into a number of quasi-independent groups of correlated amino acids termed “protein sectors.” (McLaughin 2012; Halabi 2009). For the authors, each sector is physically connected in the tertiary structure, has a distinct functional role, and constitutes an independent mode of sequence divergence in the protein family and propose that sectors represent a structural organization of proteins that reflects their evolutionary histories. This dichotomy led to some interesting results in drug design (Novinec 2014)

Communities or sectors or any set defined from an evolutionary analysis, a functional analysis, a dynamics analysis, and a purely structural analysis can lead to understanding, and few definitions are mutually exclusive. Sets are therefore a universal mechanism to cover user-defined schemes for biological molecules analyses and properties.

## Structural domains, subdomains and existing nomenclatures

**Element #3b: Structural domains (as well as evolutionary defined domains) should be defined as system sets and subsets.**

*Subsetting can also be seen as a mechanism to allow superseding of default domain definitions.*

As mentioned earlier, a set can be seen as a structural definition and as an annotation. Domains whether evolutionary defined (conserved across species), or structurally defined (conserved across protein structures) have this dual face: one can ask for the set of domains named/defined as “DNA-clamp” or ask for a particular domain “DNA-clamp” in a given chain.

With CDDs (Marchler-Bauer 2015), it is the case; one can see them as a set (all known sequences, including sequences of known structures). I cannot however in a chain, while looking at the structure in Cn3D, use the information as a subset. There, domains are structural, yet these definitions are not named and are not available system-wide. Sets such as domains should be systemic, available system wide as annotations and sets: In a chain associated with particular sequence/structure, system wide as a set of all sequences/structures. I should be able to select all

To perform comparative structural analyses between molecules and assemblies alike, we need structural domains such as SCOP (Lo Conte 2000, Chandonia 2017), CATH (Dawson 2017), ECOD (Cheng 2014) or NCBI structural domains if they were given a nomenclature (Madej 1996, 2014), common to these tertiary or quaternary molecular systems, not only used in the core computations but also available at the user discretion, in the same way CDDs are. A mapping between structural and evolutionary defined domains should be available, although delineation is a real issue but we should come to terms with all these definitions to truly integrate structure and sequence domain information.

### Superseding definitions

The issue of bridging with widely used definitions of structural domains in the literature is critical. Whatever definition is used as a system-wide default, widely used definitions in the community should either be available system-wide or a user may be able to introduce them (or their own) as user-defined sets, if these definitions are preferred for a user-defined analysis.

This is an issue of generality: the ability to define sets of entities in the system and name them for use in analysis. The also implies the computation and visualization of some properties at the level of user defined sets.

## Visualization and Geometrical elements

**Element #4: Allow flexible geometrical analysis using molecular features as well as abstract geometrical constructs such as points, lines/vectors, planes, spheres/ellipses, [possibly boxes], fitting selected molecular subsets.**

*Provide H-bond and non-bonds entities within boundaries and between interacting entities/sets.*

*Allow various level of granularity with drill down and roll up. (This is important, for example centroids such as center of Gravity can be defined systematically at the residue, set, domain, or molecule level)*

Geometrical constructs such as best-fit planes/axes/centroids associated with molecules and substructures should be definable by users for analysis. Some can be pre-computed. Straightforward geometrical properties such as distances between centroids, axes, angles between axes and planes should be available; Frames of Reference of molecular assemblies and subsets should be available/visualizable, especially when comparing assemblies or parts.

### Entities, Interfaces between entities

As mentioned before, technically and ideally any sets of atoms, residues, molecules could become a named entity to be analyzed in terms of its internal interactions as well as its (external) interactions with other entities. In pre-computing interactions, entities are naturally defined as domains, however the ability for the user to define its own entities could be possible in the interactive/post processing analyses. Arbitrary sets can solve the generality problem. This means user can define pick, list or read-in lists of molecules, residues, atoms. All this should be available through Elements #1- #3.

In 3D, many dimeric interfaces are possible, even between similar domains. On the other hand, some assemblies show some preferences. Clearly the ability to compare and differentiate and classify such dimeric interactions within interacting systems is a must. In the example of lectins (Brinda 2005 - see Figure https://www.ncbi.nlm.nih.gov/pmc/articles/PMC1237133/figure/F1/), the three-dimensional structures of the dimeric interfaces of the legume lectins (II, X4, X3, X1 and X2), galectins (galectin-1, galectin-2 and galectin-7) and pentraxins (SAP and CRP) are shown in ribbon representation with the monomers differentiated using different colors. Similarities and differences can be shown/computed both qualitatively with graphics and quantitatively with distance/orientations, sequence-structure interactions, and differences of interactions. In this example, a few centroids/axes/planes and associated variables can describe the overall problem geometrically and simplify the overall analysis, in conjunction with purely molecular interactions responsible for the variant geometries (intermolecular H-bonds, electrostatic, hydrophobic, surface interactions, salt bridges etc.).

Lectins can also be monomeric (arcelin-5, galectin-3, Charcot–Leyden protein, calnexin and calreticulin) and share a jelly-roll tertiary structure. This raises questions about what sequence/structures will dimerize. This is a central question in conformational diseases.

All assemblies can be studied at the geometrical level, as n-body problems to **spot where to look for differences,** using for example at most n(n+1)/2 distances. With very large assemblies one can limit a geometrical relationship table for example to nearest neighbors, i.e. molecules in contact, to reduce the size to a manageable dimension. In all cases zooming on a subset of interacting partners will likely be sought in **getting into details as a second step**. That can be done selectively for parts, iteratively, interactively and/or through post processing with specific molecular variables and specific user defined pseudo-molecular variables (geometrical features, centroids, rotations-translations, symmetry operations, etc.). Any level of sophistication can be envisioned: geometric, residue interactions, contacts, salt bridges, H-bonds, solvent accessibility, self-association, evolution/conservation, energy, dynamics etc.

The list of molecular properties can be large and related to the subjectivity of users/scientists. That means a choice should be made for important properties of general interest. Users ideally could add others. There are ways to do that (for discussion).

## Generalized Visualization

Visualization of molecular structure has been associated almost exclusively with 3D Graphics, and we are still using this paradigm. It is only with the first structural homology programs, and Cn3D in particular, that visualization has taken a dual aspect of sequence and structure. Cartoons – used in almost any biological paper has always been anecdotal, although it is present in each and every page of MMDB “Structure summary”. While it is a summary it could be a lot more and be dynamically linked to both sequence and structure windows. Finally, Tables have just recently been discussed to represent and control the visualization of elements of an assembly, with names. While Tables are discussed, it is worth considering them as a 4^th^ type of visualization as in any data analysis field, and in our case, think of a dynamic link to structure. (There are precedents to learn from). So to study quaternary structure one may also use 4 visualization “dimensions”.

**Element #5: Allow Visualization of molecular entities and their interactions with synchronized windows/workspaces: 3D graphics, sequence browser, 2D representations and Tables. Selecting/Highlighting any part on any of these workspaces should be reflected on any other.**

## 3D Molecular Graphics

**Element #6: The same (perceived) speed should be available to view a molecular assembly containing tens of thousands of atoms and small molecules. Widely used graphical representations such as Ribbons, CPKs, surfaces, etc. should be available. Default representations should be set according to the context of molecular assemblies depicted for maximum clarity.**

*Export of graphical snapshots as well as coordinates and topological definitions of selected assemblies, molecules and sets should be available, in widely used formats.*

Assemblies can be compared with 3D graphics, using:

- **Superimposed** assemblies using various visual representations already commonly used as well as some showing where differences occur.
- **Differential visualization** using color coding, displacement vectors of parts (loops or entire domains or subdomains, at some visually manageable/meaningful level), solid sausage (as used sometimes to represent flexibility/variability in NMR structures), etc.
- **Parallel windows** is another way to show differences, where each assembly is in a separate window in the same frame of reference, and orientation, where similar colors are used for common and unchanged parts vs. what is different and/or has moved, synchronized on color, orientation … to visualize in parallel a couple of assemblies or subsets. The same holds for a synchronized sequence window as well. Naturally switching back and forth between superimposed and split parallel windows should be possible, maybe even using them together.

### Example

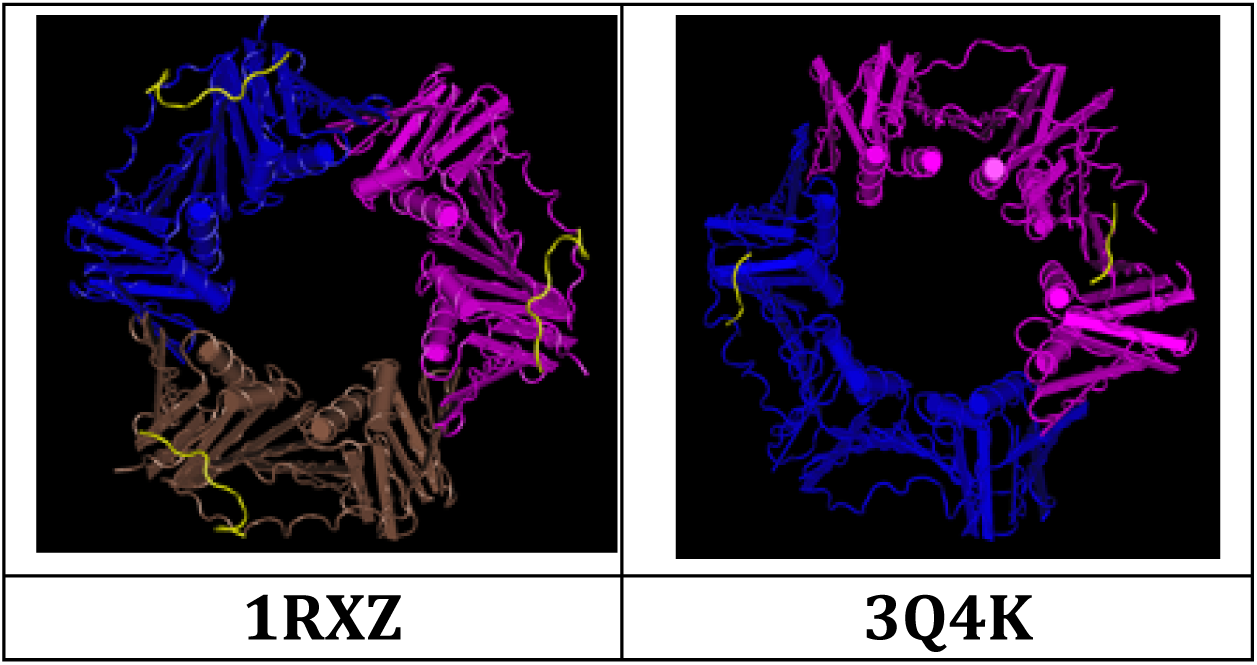

In this example, using Cn3D, one can see differences between a dimer and trimer forming a quaternary assembly as a similar toroid ring. In terms of domains these toroids are superimposable, yet they retain divalent vs. trivalent binding accordingly to their quaternary structure, even if binding sites at the interface between domains are similar and superimposable. A superimposed view as well as a dual visualization with parallelized side-by-side windows in a common frame of reference can give a clearer picture of differences than a single superimposed one. It can be a useful addition to classical superimposed views.

The important graphics feature is to be able to **spot and measure similarities AND differences in structure/conformation and correlate them with changes in sequence, function (ligand binding, dimerization …), and/or conditions.**

## 1D Sequence

**Element #7 – Sequence browser: An (extended) Sequence browser for proteins, RNA, DNA and small molecules should be available with as many tracks as needed but always remain manageable whether one looks at interacting protein sequences, experimental data, protein-RNA, protein-RNA and small molecules interactions at any the level required.**

*Tracks associated with secondary structures, user defined sets, and mutations should be available*

*Visualization of full sequences alignments or choice of a reference sequence and mutations tracks should be possible.*

*Links to corresponding databases and reference data sets, such as the human (or other reference) genome should be automatic.*

Assemblies can be compared at the sequence level in sync with 3D graphics (a la Cn3D sequence window). Sequence should be augmented with secondary structure and other tracks as needed to annotate sequence with structural or functional information obtained from 3D structure analyses. Representations on sequence beyond color should be sought out. Tracks are a simple way to look at parallel annotations, and allow correlations to emerge.

Cn3D has colorings based on basic properties that should also be used, naturally, especially in sync with 3D graphics. A good number of genomic information can be mapped from DNA to protein tracks. A straightforward example is the display of exons boundaries on protein sequence and 3D structure, or the delineation of domains and sets, as done today using color. A variation browser exists now at NCBI, and its functionality could be embedded as well, especially to map onto 3D structure mutational information.

### Example

Beta-2-microglobulin is an Immunoglobulin-like beta barrel, with 7 beta strands in a Greek key topology. Structural annotations, including secondary structure and loops information can be “tracked” and compared in tracks as well as in 3D with annotations that can be shared and visualized across sequence, annotation tracks, and 3D. (See Karamanos 2014 – Figure https://www.ncbi.nlm.nih.gov/pmc/articles/PMC4104025/figure/fig1/)

Bringing annotations together in sequence and structure, showing significant mutations in a track, and immediately locating mutations and changes in secondary structure, showing their effect in 3D, such as contact changes would be very useful. Important information noticed in 3D can be captured as an annotation/track that can then be saved and redisplayed at will, even relating the structure later to yet another structure for comparison purposes.

**Sequence is a special “dimension”** as it is rooted in genes and genomes. It is a potential windfall, yet unused! Opening up that link, that channel to navigate from protein structure to sequence and all genomic information/annotations is also enabling navigation in the other direction: opening up all structural annotations to genomic browsers. This can be a significant progress for multidisciplinary research in biology.

We could effectively go much further into a multilayered **protein sequence browser,** with layers of annotations, related to sequence and shown as appropriate tracks. Annotations of the sequence based on structure would enhance current capabilities tremendously. User annotations, through user annotation tracks that could be saved by the user in his project would be a plus. Mapping sequence (mutations) and annotations on structure as demanded and where it makes sense (for example exons, or even a label on function) would also be useful

### Remark

**What makes the strength of the whole genomics field is the systematic use of a common and unique one-dimensional coordinate system, i.e. the genome, accessible through a genome browser where annotations are represented in parallel tracks**. The multiple dimensions attached to genomic sequence such as expression, epigenetic marks and states such as methylation etc. can all be represented as tracks, and correlations such as methylation vs. expression emerge naturally. Multiple tracks can be seen as a profile. In providing a protein sequence browser in sync with structure and functional properties in sync with genomic properties would bring structure and structural interactions a step closer to mainstream biology. It may be a good time to **start conceiving a sequence browser and a bridge to genomes and genomic information.**

As a side benefit, and there should be many more, one would also know where proteins in an assembly are located in a genome, and this can be useful information, especially when assemblies compared have some functional/dysfunctional relevance, especially when more and more differential gene expression may become available.

Bringing genes and proteins within a common browser may be attractive enough to scientists currently looking at the genes that may be interested to look at proteins tracks on this particular platform instead of using multiple expert links as currently. Naturally the level of details is not the same for a structural biologist and a bench biologist. So, there should be different levels of tracks/annotations, and the ability to drill-down and roll-up among various levels of complexity may be a way to address the problem of universality of a browser.

A list of important system annotations of general interest should be defined. We already mentioned some such as secondary structure, domains, structurally conserved regions, loops, ligand binding sites, conformational variability and conformation triggers such as proline cis-trans isomerization site, beta bulges with anti-aggregation function, etc. 3-10 helices, key structurally/functionally conserved residues, conformational antibody binding sites, binding sites of all kinds. 3D domain or residue interaction networks, H-bond networks at any level of detail: residue, domain, subdomain, arbitrary sets; 3D H-bond Networks are important information to consider. Protein-protein Interactions may also be represented in appropriate representations. Protein (active sites) – small molecule ligands/drugs represent an area that is covered by many tools/databases, but that could be very useful especially in comparing proteins and protein assemblies.

**Element #7b - Allow alternate forms of visualization of interacting parts of a molecular assembly (interactomes), at various levels of details (drilling down from proteins, domains, sets, residues)**

*For example, a circular sequence browser with all interacting parts linkable with interactions (as often used in genomics (CIRCOS-like) is an option).*

In most cases, a single view or representation will not get us to deep understanding and “aha moments”, but the use of multiple simultaneous views at the desired or multiple hierarchical levels (molecules, domains, subsets, residues, atoms) may. A generalized molecular viewer should be about visualization of interacting structural domains, drilling down to local interactions in sequence and structure, and the ability to extract/abstract, represent and correlate key information.

## 2D Cartoons

**Element #8: Allow common and new 2D depictions that represent the assemblies under investigation, the sets selected, domains, secondary structures and their interactions, at any level of detail required. A synchronous view of 2D and 3D should reflect selections in either one or the other window**

*Examples of 2D representations: a cartoon of an assembly, a 2D small molecule representation, a H-bond networks between selected sets.*

### Current (2D) Cartoons of interacting parts

The example below, obtained from a VAST+, shows a comparison between 2 assemblies (3CW1 and 3S6N). It could be improved by offering a parallel view with all matching entities with the same position, colors and representation, to easily spot what is common and what is different.

Parallels with canonical domains such as CDDs or canonical assemblies would be informative rather than merged, as is currently the case. Also, users should also be able to show more or less details, and their selection could be reflected in the 3D graphics window and vice versa (maybe greying the undisplayed entities). In other words turn the 2D as a selection panel as the same time as a representation.

#### A cartoon as a graphical abstract

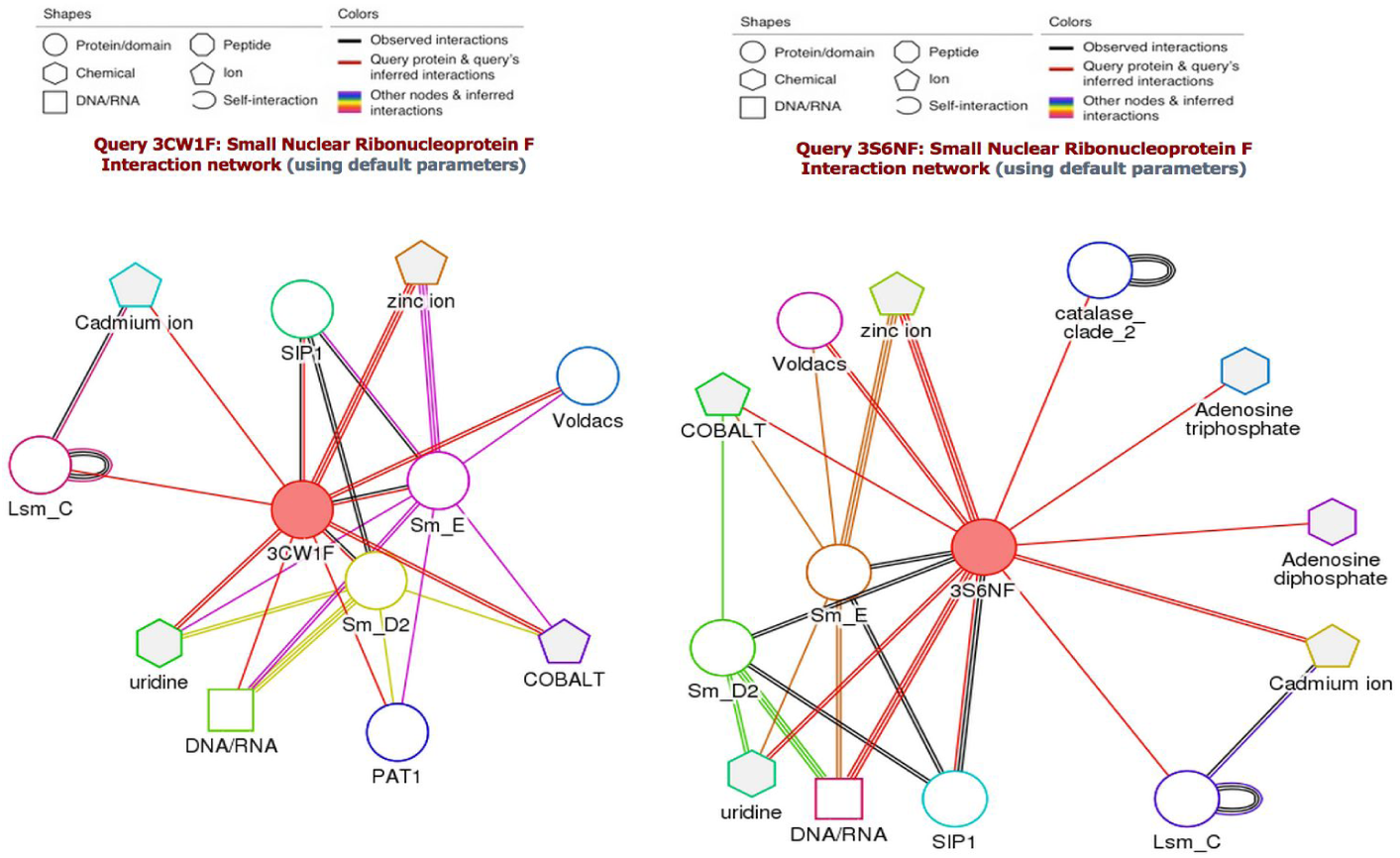

Although a 2D cartoon representation mapping the assembly organization is useful, it can difficult for general use with large assemblies, unless we simplify it. In the case of toroids as in the example above and below, it is easily doable and useful to map 3D information on a 2D cartoon, something like an assembly topology. In fact an example, we can see such cartoons used as graphical abstracts in publications (see in this instance Zhang 2011). One can effectively think of a cartoon or abstract depiction, literally and graphically, summarizing here a story of the differences between the assemblies compared.

- The assembly in file (PDB) 3S6N is the Gemin2-SMN-Sm pentamer leading, by association with a dimer of Sm proteins B-D3 form, to a Sm toroid heptamer. The first represents a snapshot with SMN acting as a chaperone of the Sm pentamer to assemble with the Sm dimer and form a functional Sm heptamer toroid core (D1-D2-F-E-G-D3-B-D1).
- The assembly in file (PDB) 3CW1 is the Sm heptamer binding U1 RNA

There is a lot to learn in analyzing and comparing assemblies, especially when they can be looked at states (snapshots) in a dynamical process. Many more assemblies in the future are expected to provide snapshots in dynamical biological events, and information can be integrated and mixed between EM and XRay.

2D abstracts can provide a reduction in complexity to illustrate assemblies and processes, as much as interactive views in parallel to 3D graphics and sequence.

In terms of existing capabilities, 2D Topologies of small molecule ligands are already cartoon depictions at the atom or fragment level. These representations can be synchronized to highlight atoms or fragments in 2D/3D and binding interactions.

We have seen earlier the example of community analyses where 2D cartoons can be used in sync with 3D to analyze molecules (or assemblies) with a user defined parts. When comparing two or more assemblies one can follow/spot differences with a 3D representation coupled with a 2D representation, sequence and naturally numerical properties. All of which can be simultaneous highlighted and colored coded according to quantitative or qualitative properties.

## Tables

**Element #9: Allow Tabulation of sets at any level of detail with tables for sets members and their properties and multiple entry tables for interactions. Synchronization with 1D, 2D, 3D views should allow selection from any window and simultaneous highlighting in the others.**

*This means a Table is a representation of properties as much as control dashboard for selection and highlight of entities, their interactions in appropriate workspaces, at any level of detail, with drill down as appropriate from molecules to domains, to sets to residues and atoms.*

A Table is a relational object, so the question is what entities do we want to relate? It solves how we can relate named entities: lined up in rows or columns. Entities represent molecular objects. There is a natural resolution/granularity to these objects that can be used for drilling down: its hierarchy, going from assemblies to molecules (RNA, DNA, organic small molecules essentially), [Domains, Subdomains], Residues, Atoms.

Representations of molecular objects can be visualized in 3D (molecular graphics, 3D plots), 2D (topologies small molecules, 1D (sequence or sequence based profile). A table can be a control table for all the representations in corresponding application/representation windows. These windows can communicate and be in sync on the data (selected, compared, highlighted, colored).

The most important structural properties (a list of which should be established) may need to be tabulated as well represented in 3D or 2D and linked as annotations to 1D (sequence). One can think of pre-computed properties. Also, since interactive operations may for example re-align structures, recomputed properties such as Pairwise RMS deviations of entities for a given assembly alignment, distances between centroids etc. based on realignments, etc.

## Structural Comparisons

**Element #10: Allow flexible pairwise (and if possible multiple) structure comparisons across molecular assemblies at any level of detail.**

*Allow the transfer of subset definitions when possible between corresponding similar entities (mapping) in compared assemblies.*

*Allow superimposed as well as split windows synchronized views for individual assemblies within a common frame of reference.*

Currently VAST+ does pairwise superimpositions of assemblies (of proteins) in PDB files for sets of proteins. As discussed, it would be useful to have **superimpositions** for pairs of assemblies – from various “points of view” i.e. using various reference structures in the assemblies, down to various meaningful substructure levels for:

- **Biopolymers: proteins, RNAs, DNAs** (ssDNA, dsDNA in particular), for any set/substructure (RNA features?), down to the residue and atom level.
- **Small molecule ligands:** On small molecules, common fragments could be superimposed when common, otherwise for example Centroids such as Center of Mass and a local frame of reference (axes along moments of inertia) could be used. Choosing a set of atoms would work universally, however pre-defined organic fragments for small molecules would be of value (equivalent to residues).

Naturally some alignments can be **precomputed** and others have to be computed **on the fly,** as in Cn3D does today for a realignment of a VAST alignment through **post processing,** if a user looking at the assembly/assemblies wants to look at “another angle” of the problem and change perspective/reference, or define him/herself an “invariant set” across assemblies. It may be prohibitive to pre-compute all pairwise comparisons and it may not be all that useful. However, the ability for a user to change reference and re-compute alignments based on that reference, based on some user-imposed constraints (an angle) may be very powerful and not that compute intensive depending on restraints.

Visual and tabular information on **relative positions and orientations** of molecular entities in an assembly for a given superimposition would be useful (in short: geometrical properties such as distances between centroids, axes, displacements along and rotations relative to axes…), as a way of analyzing and comparing assemblies.

The availability of **transformation matrices** (rotation-translation) and/or rotation axis, angle, translation vector, could prove valuable. [Provision for retrieval and storage of transformation matrices (rotation-translations) between global and local frames of references associated with assemblies, molecules or arbitrary sets, to be applicable to other datasets. Or, a sort of copy/paste in a given transformed position could be useful to test hypotheses for example on dimerization. This may be seen as modeling as opposed to analysis, yet it would be useful in some cases to position a structure for comparative analyses, provided we have tools to analyze interfaces in terms of H-bonds, non-bonds, charge-charge, surface contacts, and clashes, which can eventually be further studied with other third party energy based tools].

**Element #11: Usability - Allow storage/retrieval of projects, with all annotations, if possible as a database. Eventually the analyzed assemblies stored in a project can become a seed for a real database for downstream projects (such as drug design).**

**Element #11b: Usability - Simple and unique mode of user-software interaction, systematic synchronization of highlighting, labeling, annotating between windows (and tracks) with a native hierarchical drill down.**

## The 12^th^ Element: Symmetry in Tertiary and Quaternary Structures

In quaternary structures, non-XRay crystallographic symmetry is ubiquitous: **80% of PDB biological assemblies exhibit point group symmetry** (unpublished - This number deserves confirmation). It also means we may have means to use symmetry in calculations to improve performance as well as analyze symmetry properties. This should be particularly significant with very large viral assemblies for example, that exhibit a very high order of symmetry. **Pseudo-symmetry is also clearly present in ca. 20%** of structural Superfamilies as was demonstrated recently through a census on the whole PDB (Myers-Turnbull 2014).

***The question is: can we analyze assemblies without considering symmetry?***

### Structural Properties of assemblies and domains within assemblies: Tertiary vs Quaternary structure analysis

What matters is a biological unit, whether tertiary or quaternary structure. A tertiary structure, a protein chain, can also be seen as a covalently linked assembly of protein domains. In which case linkers can now be seen as these determinants of geometrical relations between domains, developed during evolution, a whole new area to be explored systematically…

So what is technically the difference between a tertiary assembly and a quaternary assembly of interacting parts? And in terms of domains and domain interactions how does it matter?

## A few more issues for brainstorming and discussion

Many questions can be asked. Here are a few just as a seed:

- What are the main structural properties of general interest? And how to extend the repertoire of properties accessible dynamically?
- What other representations may be of use? or data management elements may be needed? Tables for example? Graphs? Else?
- How to enable widely used as well as new user defined Nomenclatures and allow them to evolve?
- What properties may relate structure to function, to known dysfunction, or to evolution?
- Etc.

This white paper originally started as a list of requirements. There is however a limit to what a list of requirements can achieve, as there are many applications to think through, some unforeseen. Twelve is an arbitrary number, the analytical process is universal and many of the data types involved are already known and reusable. The idea is to help developers think about key concepts that may be of use in assembling powerful molecular structure analysis tools. Do the twelve elements cover enough of a common ground? The list of molecular properties of interest to all can be very long. However as long it may get though, **do the Twelve Elements conceptually enable most structural analyses?** That question should be a test to think about the extensibility of the system.

Euclid’s Elements contained 13 books. Any interested developer could add his or her own 13^th^ Element, as much as amending the proposed twelve for the better. It is time to get into action in an agile and open source environment. I only hope the Elements can help structure some thinking about new software components and APIs, ultimately some well-organized 3D Molecular JavaScript libraries (not forgetting lower and higher dimensions), similar to 3D JavaScript libraries such as three.js.

## Conclusion: A few Elements in Action in current web viewers

Two years into development, web viewing using JavaScript and WebGL unveils immense capabilities. Although still in its infancy, this could signal the start of a new era for molecular graphics. It is important however that web viewers do not entrench themselves in simply rewriting 3D viewing capabilities, currently available in other environments aimed at experts in structural biology, molecular modeling, protein engineering or drug design. Developers should try to project ahead an integrated approach to visualize and analyze molecular entities and their interactions in multiple dimensions, at multiple levels of details, for diverse users, bridging 1D sequence and 3D structure representations and databases, at a minimum.

All current web viewers partially demonstrate some of the Twelve Elements:

Element #1 : Naming of molecular entities
Element #2 : Enable import of molecular structure coordinates and topologies
Element #3 : Flexible set selection of any part of molecular entities
Element #3b: Systematic Structural or Evolutionary Domains definitions as sets
Element #4 : Geometrical analysis using molecular features and geometrical constructs
Element #5 : Visualization of molecular entities and interactions in 3D, 2D and Tables
Element #6 : Speed for 3D manipulation of very large molecular assemblies
Element #7 : Sequence browser/1D Track management for proteins, RNA, DNA
Element #7b: Visualization of interacting parts of a molecular assemblies (interactomes)
Element #8 : Common and innovative 2D depictions to represent molecular assemblies
Element #9 : Tabulation of sets & sets elements and their properties at any level of detail
Element #10: Flexible pairwise (and multiple) structure comparisons
Element #11 : Usability - Storage/retrieval of projects, with all annotations
Element #11b: Usability - Simple and unique mode of user-software interaction
Element #12 : Integrated Symmetry Analysis

All viewers for example, demonstrate Elements #1 and #2. **The NGL viewer**, available in Github (https://github.com/arose/ngl), does brilliantly demonstrate Element #6, as it allows in particular, fast loading and fast 3D manipulation of very large viruses across networks. The **iCn3D viewer**, now released in its version 1.0 is operational on NCBI’s server pages, and does demonstrate *in part* Elements #3, #5, #7, #8, #10:https://www.ncbi.nlm.nih.gov/Structure/icn3d/docs/icn3d_about.html. This is only a beginning. The road ahead is still long but can be shortened if one can rely on web component libraries to grow functionality in visualization and analysis in a sustainable and scalable manner. The code is available in GitHub (https://github.com/ncbi/icn3d).

In the example used earlier in the text, iCn3D can be used to compare two molecular assemblies superimposed (a trimer of fused domain dimers PDBid:1RXZ vs. a dimer of fused domain trimers PDBid: 3Q4K), using a 2D cartoon to represent and identify molecular interactions between various chains in an assembly and highlight these interactions at the residue level in 1D sequence and 3D structure. https://www.ncbi.nlm.nih.gov/Structure/icn3d/full.html?showseq=1&show2d=1&align=1rxz,3q4k. Future Improvements should allow drilling down at the domain and secondary structure level, yet it is already quite straightforward to analyze interactions.

As agile development is becoming the norm, an idea can be tested quickly through a prototype. We have been experimenting during hackathons on how to enable information flow between a web based genome/protein sequence 1D browser and a 3D viewer: (https://github.com/NCBI-Hackathons/Structure/Visualization), mapping and visualizing SNPs from dbSNP or ClinVar databases in protein coding regions onto 3D protein structures directly from a genome browser. Beyond the ability to visualize, manipulate, compare or edit 3D molecular structures without the need for a separate app, data can be exchanged dynamically between the 3D viewer and a 1D genome browser. The web environment enables bi-directional mapping for automatic or user-controlled annotation of 1D sequences with 3D structure information, and vice versa, without expert knowledge. This opens new perspectives, as molecular properties and molecular interactions can be fingerprinted, visualized compared in both contexts, 1D genome browsers and 3D viewers, and shared: https://www.ncbi.nlm.nih.gov/Structure/icn3d/full.html?showseq=1&show2d=1&mmdbid=2ROX&command=color%20grey;%20%20select%20:119;%20style%20sidechains%20sphere;%20%20color%20red;%20%20select%20zone%20cutoff%205;%20%20color%20blue;%20%20toggle%20highlight;%20pickatom%202876;%20pickatom%201845;%20pickatom%201834;%20pickatom%201843;%20set%20surface%20molecular%20surface;%20toggle%20highlight;%20%20add%20label%20SNPT119%20%7C%20size%2018%20%7C%20color%20ffff00%20%7C%20background%20cccccc%20%7C%20type%20custom. In this simple example, Residue T119 in the wild type Transthyretin (PDBid 2ROX), a SNP location is highlighted in red. Residues and ligands in contact within 4A are colored in blue.

Sharing instantaneously annotated molecular graphics representations through the Web is one of the most amazing capabilities of 3D web viewing, and opens perspectives for collaborative research. One can potentially summarize a complete analysis and visualization through a single URL, with all needed labeling, coloring, surfacing, highlighting and commenting on sequence and structure. iCn3D has now a beta version of a **“share-link”** command which will package as a URL all necessary data and operations/commands to share a 3D/2D/1D visualization in a given state, similar in essence to Google maps that can be shared through URLs:

*If a picture is worth a thousand words, and if an interactive 3D graphics representation is worth a thousand images, then these 3D interactive shared URLs could be worth a million words. Anyone can draw his or her own conclusions on the impact of this technology.*

## Acknowledgements

This research was supported in part by the Intramural Research Program of the NIH, National Library of Medicine. Special thanks to Jiyao Wang who developed iCn3D, with whom I had the pleasure to collaborate, as well as members of NCBI’s Structure group headed by Steve Bryant, and in particular Dachuan Zhang, Aron Marchler-Bauer, Tom Madej, Christopher Lanczycki, Lewis Geer, Renata Geer, Yanli Wang.

